# Tracking of antibiotic resistance transfer and rapid plasmid evolution in a hospital setting by Nanopore sequencing

**DOI:** 10.1101/639609

**Authors:** Silke Peter, Mattia Bosio, Caspar Gross, Daniela Bezdan, Javier Gutierrez, Philipp Oberhettinger, Jan Liese, Wichard Vogel, Daniela Dörfel, Lennard Berger, Matthias Marschal, Matthias Willmann, Ivo Gut, Marta Gut, Ingo Autenrieth, Stephan Ossowski

**Author notes:** These authors contributed equally (Mattia Bosio, Caspar Gross).

## Abstract

**Background:** Infection of patients with multidrug-resistant (MDR) bacteria often leave very limited or no treatment options. The transfer of antimicrobial resistance genes (ARG) carrying plasmids between bacterial species by horizontal gene transfer represents an important mode of expansion of ARGs. Here, we evaluated the application of Nanopore sequencing technology in a hospital setting for monitoring the transfer and rapid evolution of antibiotic resistance plasmids within and across multiple species.

**Results:** In 2009 we experienced an outbreak with an extensively multidrug resistant *P. aeruginosa* harboring the carbapenemase enzyme *bla*_IMP-8_, and in 2012 the first *Citrobacter freundii* and *Citrobacter werkmanii* harboring the same enzyme were detected. Using Nanopore and Illumina sequencing we conducted a comparative analysis of all *bla*_IMP-8_ bacteria isolated in our hospital over a 6-year period (n = 54). We developed the computational platforms *pathoLogic* and *plasmIDent* for Nanopore-based characterization of clinical isolates and monitoring of ARG transfer, comprising de-novo assembly of genomes and plasmids, polishing, QC, plasmid circularization, ARG annotation, comparative genome analysis of multiple isolates and visualization of results. Using *plasmIDent* we identified a 40 kb plasmid carrying *bla*_IMP-8_ in *P. aeruginosa* and *C. freundii*, verifying that plasmid transfer had occurred. Within *C. freundii* the plasmid underwent further evolution and plasmid fusion, resulting in a 164 kb mega-plasmid, which was transferred to *C. werkmanii*. Moreover, multiple rearrangements of the multidrug resistance gene cassette were detected in *P. aeruginosa*, including deletions and translocations of complete ARGs.

**Conclusion:** Plasmid transfer, plasmid fusion and rearrangement of the multidrug resistance gene cassette mediated the rapid evolution of opportunistic pathogens in our hospital. We demonstrated the feasibility of tracking plasmid evolution dynamics and ARG transfer in clinical settings in a timely manner. The approach will allow for successful countermeasures to contain not only clonal, but also plasmid mediated outbreaks.

## Background

The increase of multi-drug resistant (MDR) bacteria has led organizations such as the World Health organization (WHO) and the US Centers for Disease Control and Prevention (CDC) to categorize MDR bacteria a major public health problem [1]. Infection or colonization of patients with MDR bacteria often leave only very limited or even no treatment options, thus posing a potentially life-threatening risk to the individual patient in particular on intensive care units [2, 3]. In addition, infection control measures to prevent spreading are required, resulting in increased efforts in patient care and costs for healthcare providers and the public healthcare systems [4, 5]. Although action is needed on different national and international levels, understanding colonization, infection and transmission routes of these MDR resistant bacteria in the local hospital setting represent a crucial initial step towards implementation of harmonized, successful strategies to combat infections caused by MDR bacteria [1, 4].

Next-generation sequencing (NGS) has become widely available and was used successfully to resolve outbreaks and determine transmission routes (e.g. reviewed in [6]). However, not only the clonal transmission of MDR bacteria, but also the spread of multidrug resistance plasmids by horizontal gene transfer between different bacterial species represents an important mode of expansion of antimicrobial resistance genes [7]. Although multidrug resistance plasmids and plasmid transfer have been studied in hospital settings, their interrogation is not part of routine infection control practice. Moreover, plasmid characterization and comparison based on short-read sequences is error prone and unreliable particularly when larger plasmids (>50 kb) are involved [8], while long-read de novo assembly based plasmid analysis is currently limited to large centers with access to Pacific Biosystems (PacBio) Sequel sequencers (e. g. [9, 10]). Recently, the MinION long-read sequencer (Oxford Nanopore Technologies, ONT) has become more widely available, facilitating fast and inexpensive analysis of multidrug resistance plasmids, horizontal gene transfer and evolution of plasmid-born antimicrobial resistance (AMR) [11, 12]. Thus, the technology is potentially suitable for application within the hospital setting. In recent publications, Dong et al examined the microevolution of KPC carbapenemase plasmids in three clinical isolates applying Nanopore technology [13], while Lemon et al optimized the Nanopore sequencing laboratory workflow and analyzed plasmids from three clinical isolates [11]. Long-read sequences substantially increase the contiguity of de novo assemblies by spanning repeat regions, resulting in finished microbial genome and plasmid assemblies [14]. However, due to the high error rates of Nanopore sequencing, hybrid assemblers such as hybridSPAdes [15] and Unicycler [16] combine long and short reads to achieve a high base level accuracy needed for the correct identification of AMR related genes and variants. In the present study, we aimed to evaluate the application of Nanopore sequencing technology in a hospital setting and to demonstrate the feasibility of monitoring transfer and rapid evolution of antibiotic resistance plasmids within and across multiple species.

Starting in 2009 we experienced an outbreak with an extensively multidrug resistant *P. aeruginosa* clone in our hospital [17]. The strain harbored a carbapenemase enzyme (*bla*_IMP-8_), which renders all beta-lactams resistant, including carbapenems, an antibiotic class of last resort. Extensive infectious disease interventions and the establishment of a rectal screening program to identify colonized patients led to a reduction of cases. However, in March 2012 we detected the first *Citrobacter freundii* harboring the same enzyme *bla*_IMP-8_ carbapenemase [18], approximately 2.5 years after the first *P. aeruginosa bla*_IMP-8_ had been detected. Shortly after, the carbapenemase was detected in *Citrobacter werkmanii*. Since this enzyme is rarely encountered in Europe, we hypothesized, that horizontal gene transfer between the different Gram-negative bacteria had occurred within our hospital. Therefore, we conducted a sequencing study including all multidrug resistant bacteria harboring the enzyme *bla*_IMP-8_ isolated in our hospital over a 6-year period including patient and environmental isolates. We developed and established a bioinformatics pipeline in order i) to determine the sequence of the *bla*_IMP-8_ harboring plasmid and characterize all contained AMR genes ii) to identify potential transmission events of the plasmid between species and iii) to characterize the evolutionary dynamics of the plasmid.

## Results

### Comprehensive analysis platform for antibiotic resistance gene carrying plasmids

We have developed a comprehensive computational platform for the genomic analysis of clinical isolates and the monitoring of antibiotic resistance gene transfer. *PathoLogic* comprises of a hybrid de-novo assembly pipeline generating finished genomes and plasmids, genome polishing, quality control, annotation, comparative genome analysis of multiple isolates, as well as visualization of results (Fig 1). Furthermore, *pathoLogic* integrates the *plasmIDent* method, which confirms the circularity of putative plasmids by ring-closure using long reads, performs AMR gene annotation, calculates various characteristic plasmid properties, and creates a circular visualization of the annotated plasmid. Finally, the sequences of plasmids from multiple isolates of the same or different species are compared in order to identify horizontal gene transfers, structural variations (e.g. AMR gene presence/absence) and point mutations, which can further be utilized for phylogenetic or transmission analysis. *PathoLogic*, *plasmIDent* and a graphical user interface (GUI) are freely available on github (*plasmIDent* pipeline: https://github.com/imgag/plasmIDent, pathoLogic pipeline: https://github.com/imgag/pathoLogic).

**Figure 1.**
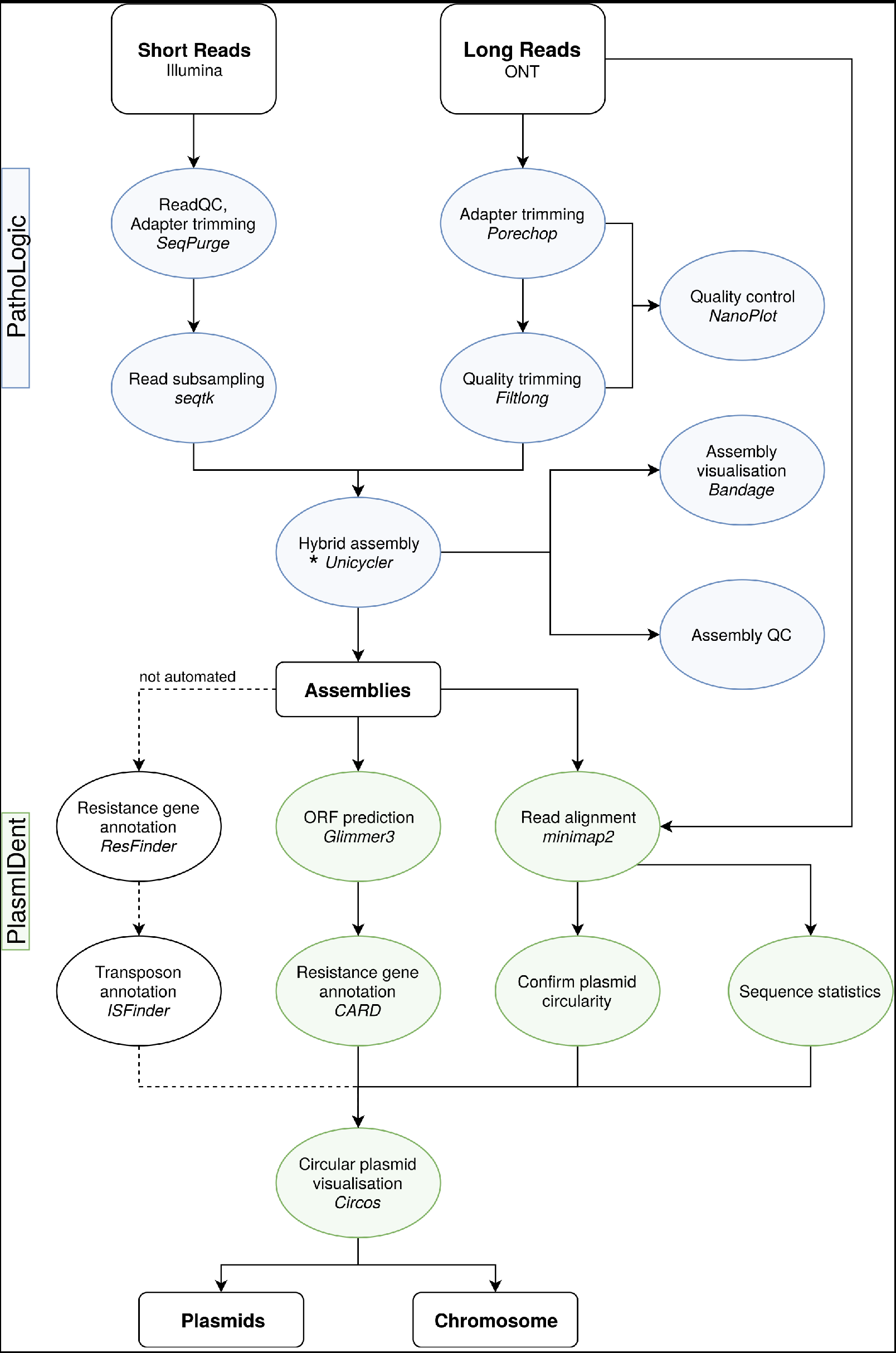
Schematic diagram of the data analysis workflow used in this study. The *pathoLogic* platform was created using the *Nextflow* [19] environment to chain different tools and scripts, represented here as circular nodes. Connecting lines indicate data flow between the separate processes, dashed lines show tools that are not directly included in the pipeline and need manual data handling. In *pathoLogic*, the assembly step (*) can be performed by *Unicycler* [16], *Canu* [20], *miniasm* [21], *hybridSPAdes* [15] or *flye* [22, 23].

### Characterization of study isolates

In our study, we included all *bla*_IMP-8_ AMR gene positive strains isolated in our hospital from patients or patient-related environmental water sources in the haemato-oncology department over a 6 years’ period (n=54). This also comprised the previously reported *P. aeruginosa* outbreak clones (n=34) [17] and one *C. freundii bla*_IMP-8_ isolate [18], for which Illumina short-read data is available (Tab. S1). In order to obtain finished genomes and circularized plasmids long read Nanopore sequencing was conducted of all *Citrobacter freundii* (n=8) and *Citrobacter werkmanii* (n=1) isolates and selected *P. aeruginosa* isolates (n=5) representing different time points (Tab. S2). Applying the *pathoLogic* pipeline described above enabled us to generate high quality genomes for all samples. In 5 of the 14 samples we were able to generate a single circular chromosome along with several circular plasmids (Tab. S3). All other assemblies also have few, large contigs, as indicated by a high NG75 values. Samples with a lower depth of coverage of Nanopore reads (e.g. isolate 9_E_CF) also resulted in more fragmented assemblies.

### Plasmid content and phylogeny of the study isolates

Within the first 2.5 years we only observed *P. aeruginosa bla*_IMP-8_ involving colonization or infection of 26 patients, before the first detection of *C. freundii* and *C. werkmanii* carrying *bla*_IMP-8_ as shown in the timeline (Fig. 2A). The plasmids with relevance to the dynamics of the *bla*_IMP-8_ plasmid evolution are displayed in figure 2B. The complete plasmid content of all isolates is summarized in table S4.

**Figure 2.**
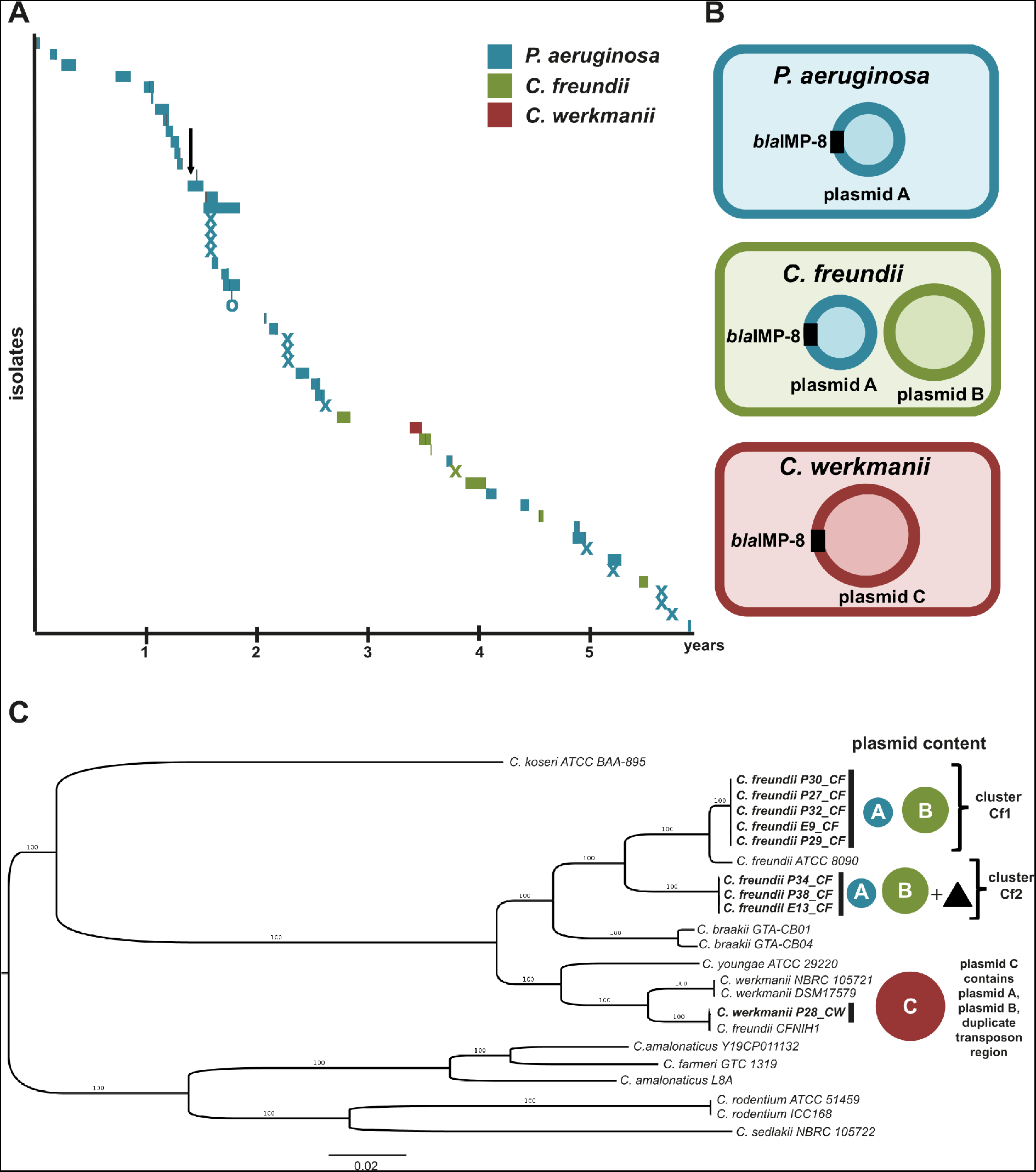
**A.** Timeline of isolation of *bla*_IMP-8_ Gram-negatives in the hemato-oncology department over 6 years. “P” indicates an isolate from a patient. For patient isolates the length of hospital stay is displayed. Patient 21 was seen only in the outpatient department, marked with “o”. Environment isolates are named with an “E”. The date of isolation is shown as an “x”. The introduction of a rectal screening program is marked with a black arrow. PA: *P. aeruginosa*, CF: *Citrobacter freundii*, CW: *Citrobacter werkmanii*. **B.** Overview of plasmids with relevance to the evolution of the *bla*_IMP-8_ plasmid found in *P. aeruginosa*, *C. freundii* and *C. werkmanii*. **C.** Maximum likelihood phylogeny of *Citrobacter* species included in the study (n=9). The *Citrobacter freundii* strains formed two clusters Cf1 (n=5) and Cf2 (n=3). Strains of cluster Cf2 harbored chromosomal transposon region (black triangle) homologous to the regions of plasmid C. *C. werkmanii* clustered with the respective reference strains NBRC105721 and DSM17579. The scale bar shows the expected number of nucleotide changes per site.

In *P. aeruginosa* isolates, we detected a 40 kb plasmid carrying the *bla*_IMP-8_ gene (plasmid A, blue). In *C. freundii* isolates the same *bla*_IMP-8_ plasmid A was found in addition to an 88 kb plasmid (plasmid B, green) without a carbapenemase gene. Surprisingly, in the *C. werkmanii* isolate a large 164 kb plasmid harboring the *bla*_IMP-8_ gene was detected (plasmid C, red) without any evidence for the presence of the plasmids A or B. The structures and circular nature of the three plasmids were confirmed by remapping the long-read sequences resulting in continuous read coverage along the plasmids without breakpoints.

Phylogenetic analysis showed that all *P. aeruginosa* strains are closely related and belong to a single cluster indicating clonal spread (Fig. S1). In contrast, the maximum likelihood phylogeny of the *Citrobacter* isolates revealed a phylogenetically more diverse picture (Fig 2C). The *C. freundii* isolates formed two clusters Cf1 (n=5) and Cf2 (n=3), which were clearly distinct (Fig. 2C). Both clusters contained the plasmids A and B. Isolates of the cluster Cf2 contained an additional plasmid G (Tab. S4) and a transposon region localized on the chromosome absent in cluster Cf1, which is further described below. The *C. werkmanii* isolate clustered with two reference strains for *C. werkmanii* NBRC105721 and DSM17579, as expected.

### Comparative genomic analysis and annotation of plasmids

Next, we performed multiple sequence alignment of the generated reference sequences of plasmids A, B and C (Fig. 3). To better understand the chronological order of the horizontal gene transfer (HGT) and fusion events we first performed an in-depth annotation of plasmid features, including antimicrobial resistance genes, transposons, origin of replication and GC-content (Fig. 3). Plasmid A, which contains the *bla*_IMP-8_ gene, had an average GC content of 59% (Fig. 3 green inner circle). The *bla*_IMP-8_ was located on a class 1 integron together with eight additional antimicrobial resistance genes (Fig. 3 bottom). This resistance gene cassette had a lower GC content compared to the other parts of plasmid A, indicating that this part might have been acquired from a different organism. Plasmid B had a size of approx. 88 kb, a substantially lower GC content (< 50 %) when compared to plasmid A, and lacked the *bla*_IMP-8_ integron. The largest plasmid C with a size of 164 kb comprised of plasmid A including the antimicrobial resistance gene cassette and plasmid B, as well as two large stretches of transposase elements Tn3 and IS6. Therefore, plasmid C resulted most likely from a fusion of plasmids A and B. The two fusion regions between plasmids A and B harbored a duplicated region (marked in figure 3 with a black arrow) composed of one transposon Tn3, three IS6 elements and several AMR genes. Two additional Tn3 copies interspersed by additional AMR genes extended one of the fusion regions.

**Figure 3.**
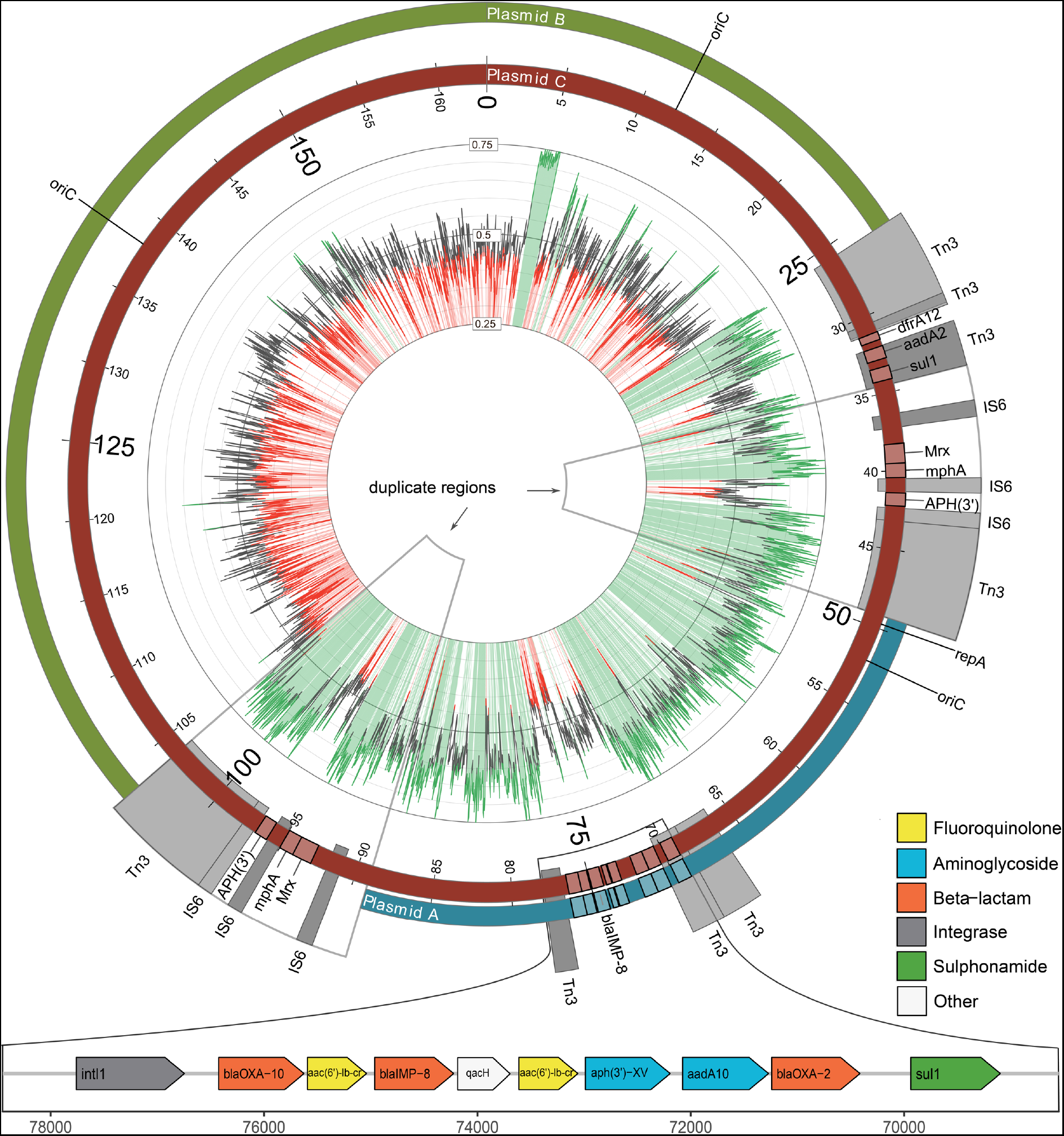
Detailed alignment of plasmid A, B and C. Plasmid A (blue, outer circle) harbored a multidrug resistance cassette including *bla*_IMP-8_ on a class 1 integron and shows high GC content (inner circle, green). Plasmid B (green, outer circle) harbored no *bla*_IMP-8_ resistance gene and shows lower GC content (red). Plasmid C (red)was the largest and comprised plasmid A, plasmid B, a duplicated transposon region (black arrows) and a unique extension by two Tn3 elements in one fusion region. The class 1 integron consists of 9 AMR genes including aminoglycosides, beta-lactams, sulphonamides and fluoroquinolones. Additional AMR and a mercury resistance operon [24] are found within the duplicate transposon region.

### Plasmid content of isolates and plasmid fusion

In order to determine the plasmid content of all studied isolates we realigned the Illumina short-read sequences using the assembled plasmid C as a reference, which comprises the sequences of plasmids A and B and the duplicate transposon region (Fig. 3). The coverage for each strain is displayed in Figure 4. All *P. aeruginosa* isolates contained only plasmid A, not plasmid B or C. Sequencing reads of *P. aeruginosa* that mapped to a small section of the transposon region most likely originate from the chromosome. The picture is more complex for the *Citrobacter* species, which could be divided into three groups. The *C. werkmanii* (28_P_CW) strain contained the complete plasmid C, which was homogenously covered. The *C. freundii* isolates formed two groups, one group with Plasmid A and B and the second group containing plasmid A and B as well as coverage on the transposon regions. The two groups are identical with the clusters Cf1 and Cf2 distinguished by phylogenetic analysis of the chromosomes.

**Figure 4.**
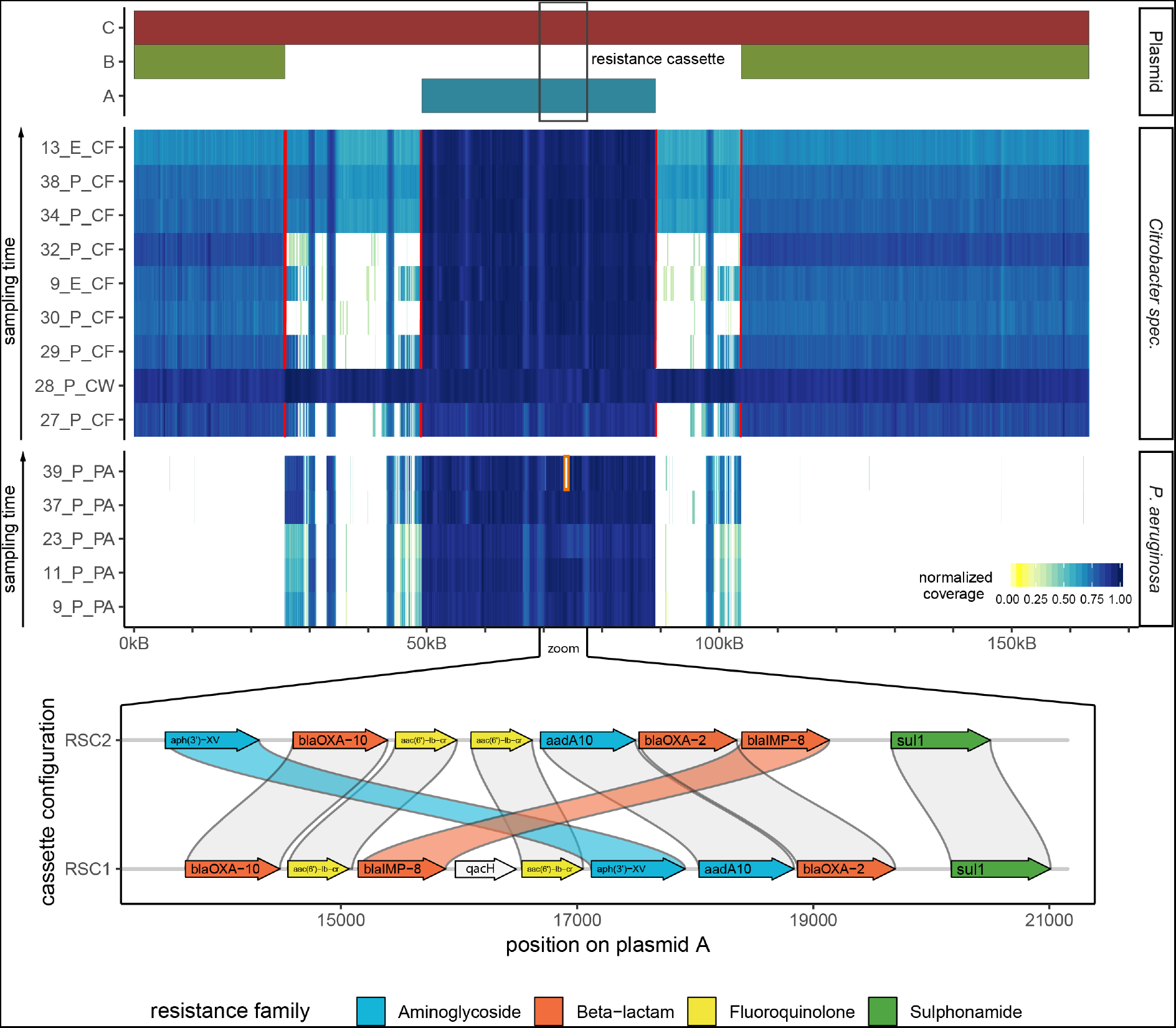
**A.** Coverage plot based on short-read Illumina data mapped against the reference plasmid C found in *Citrobacter werkmanii*. Plasmid C (red bar) comprises of the sequences of plasmids A (blue bar) and B (green bar) fused by two transposon-rich regions. Red lines indicate breakpoints, which are characterized by the absence of reads spanning the breakpoint. White areas indicate the absence of coverage and hence absence of sequence in a given isolate, which in some cases could be the result of a deletion event or indicate boundaries between scaffolds. **B.** Comparison of the resistance gene cassette configuration of RSC1 and RSC2, showing putative AMR gene translocation events and a deletion of *qacH*.

We further investigated the read coverage distribution for the *C. freundii* isolates to determine if they harbor only copies of plasmids A and B, or a combination of copies of A, B and C. No continuous short or long reads could be detected spanning the gap between the plasmid A or B sequence and the transposon regions in either of the two *C. freundii* clusters (Fig. 4, red lines indicating the breakpoints, Fig. S2 showing alignments to one example breakpoint), suggesting that the short reads mapping between A and B originate from a chromosomal integration of the transposon elements. Annotation of the assembled chromosomes of *C. freundii* and *C. werkmanii* isolates confirmed that cluster Cf2 contains the transposon sequence within the chromosomal scaffold, while cluster Cf1 and *C. werkmanii* do not (Fig. S3). We conclude that both Cf1 and Cf2 harbor only copies of plasmids A and B, but not of plasmid C, and that Cf2 harbors a copy of the transposon region in the chromosome.

Only isolates of cluster Cf2 show a complete ‘smear’ in the coverage plot across the whole transposon region (13_E_CF, 34_P_CF, 38_P_CF). In isolates of cluster Cf1, however, we observe only partial coverage of the transposon region (Cf1.1: 3 isolates, 9_E_CF, 29_P_CF, 27_P_CF) or almost no coverage (Cf1.2: 2 isolates 32_P_CF and 30_P_CF). Interestingly, the regions distinguishing these two subclasses Cf1.1 and Cf1.2 harbor a mercury resistance operon [24] present in Cf1.1 but absent in Cf1.2. Pairwise alignment to the full genomes using the *nucmer* aligner confirmed that these genes are located on plasmid J (Tab. S4) in isolate 29_P_CF and on non-circular contigs in the other two isolates (9_E_CF, 27_P_CF) of group Cf1.1 (Fig. S3).

In summary, our phylogenetic analysis as well as the comprehensive plasmid annotations indicated that the *C. freundii* isolates in the clusters Cf1 and Cf2 originate from different clones and should be treated as separate entities in the identification of plasmid-born horizontal gene transfers.

### Deletion and transposition of AMR genes in *P. aeruginosa*

While *P. aeruginosa* isolates homogeneously only contained plasmid A, we observed that for some isolates the resistance gene cassette was not continuously covered with short reads in the reference alignment shown in figure 4 and figure S4. Using short and long read based structural variant detection methods we identified two types of re-arrangement events. First, we found various deletions of resistance genes within the resistance gene cassette in 12 strains, indicated by zero coverage (Fig. S4, white areas flanked by red brackets). Analysis of the resistance genes annotated by ResFinder or CARD on the respective plasmid scaffolds confirmed that these deletions correspond to missing AMR genes in the respective strains (Tab. S5). Furthermore, all deletions span exactly from 5’ to the 3’ end one AMR gene plus the flanking IS sequence.

Moreover, when comparing the class 1 integron region of *P. aeruginosa* isolates 37_P_PA and 39_P_PA we detected breakpoints without a corresponding drop of coverage in flanking genes, indicating translocation events. We therefore performed a multiple-sequence alignment of the resistance gene cassettes of the 5 *P. aeruginosa* isolates for which Nanopore sequences were generated, as the long-read data facilitates highest confidence assemblies. Indeed, we identified two structurally different versions of the resistance gene cassette, termed RSC1 and RSC2, the latter likely being the result of multiple transposition and deletion events (Fig. 4B). Four isolates harbored the wild-type cassette RSC1, while one isolate harbored RSC2. Finally, we aligned the short reads of all 49 *P. aeruginosa* isolates against the breakpoints distinguishing RSC1 and RSC2. We identified 21 isolates most similar to RSC1 and 9 isolates most similar to RSC2, while 10 isolates could not be uniquely assigned to one or the other, pointing to a third cassette configuration (Fig S4). Our results indicate that AMR genes on plasmids are subject to strong selective pressure and are frequently removed, likely due to the high cost of transcribing multiple resistance genes.

### Rapid plasmid-mediated adaptation: acquisition and loss of AMR genes by horizontal gene transfer and structural rearrangement events

Our findings generated multiple evidences that the rapid evolution of opportunistic pathogens in our hospital was mediated by plasmid transfer, merging and rearrangement, and evolved over multiple distinguishable stages (Fig. 5) following the most likely sequence of events:

i. Plasmid A (40 kb) harboring *bla*_IMP-8_ and multiple other AMR genes was transferred between *P. aeruginosa* and *C. freundii*. Although the direction of transfer cannot be determined with certainty, the fact that *P. aeruginosa bla*_IMP-8_ was isolated approximately 2.5 years before the first *Citrobacter bla*_IMP-8_ strain was detected, suggests a transfer from *P. aeruginosa* to *Citrobacter* species. Moreover, the higher GC content of plasmid A points towards an origin of the plasmid from a background with a high GC content such as *P. aeruginosa* (average GC content of 66%). However, an unknown intermediate host serving as a reservoir for plasmid A cannot be completely ruled out.
ii. In *C. freundii*, the plasmid underwent further evolution resulting in the fusion of the acquired plasmid A and a resident plasmid B to the mega-plasmid C ultimately recovered in *C. werkmanii*. We hypothesized that this happened by plasmid fusion, since plasmid C contains regions with genetic homology of close to 100 % across the full length of plasmid A and plasmid B. In addition, plasmid C contained transposon regions, which were also present in the chromosome of *C. freundii* cluster Cf2 strains, indicating that this organism was most likely the host of the plasmid fusion. However, a plasmid fusion in *C. werkmanii* cannot be ruled out (Fig. 5 grey area).
iii. We speculate that *C. freundii* Cf2 strains ‘distributed’ plasmid A to *C. freundii* Cf1 and plasmid C to *C. werkmanii*. However, it is also possible that Cf1 and Cf2 independently acquired plasmid A from *P. aeruginosa* or that Cf2 acquired plasmid A from Cf1. Although less likely, the plasmid fusion resulting in plasmid C might have occurred in *C. werkmanii* after independent transfer of plasmids A and B from any of the other three bacteria. However, *C. werkmanii* is also lacking a copy of the transposable element region in its chromosome, which is present in cluster Cf2, making a fusion in *C. werkmanii* highly unlikely (Fig. S3). Figure 5 depicts all possible trajectories of the adaptation processes mediated by plasmid HGT leading to three bacterial species and four clones with multi-antibiotic resistances in a single hospital within a few years.
iv. In parallel, in *P. aeruginosa* the class 1 integron harboring the antimicrobial resistance genes including *bla*_IMP-8_ underwent various re-arrangements such as deletions and transpositions of AMR genes. In 12 of the *P. aeruginosa* isolates one or more AMR genes were lost (Tab. S5) and at least 9 strains show a transposition of two AMR genes within the integron (Fig. S4).

**Figure 5.**
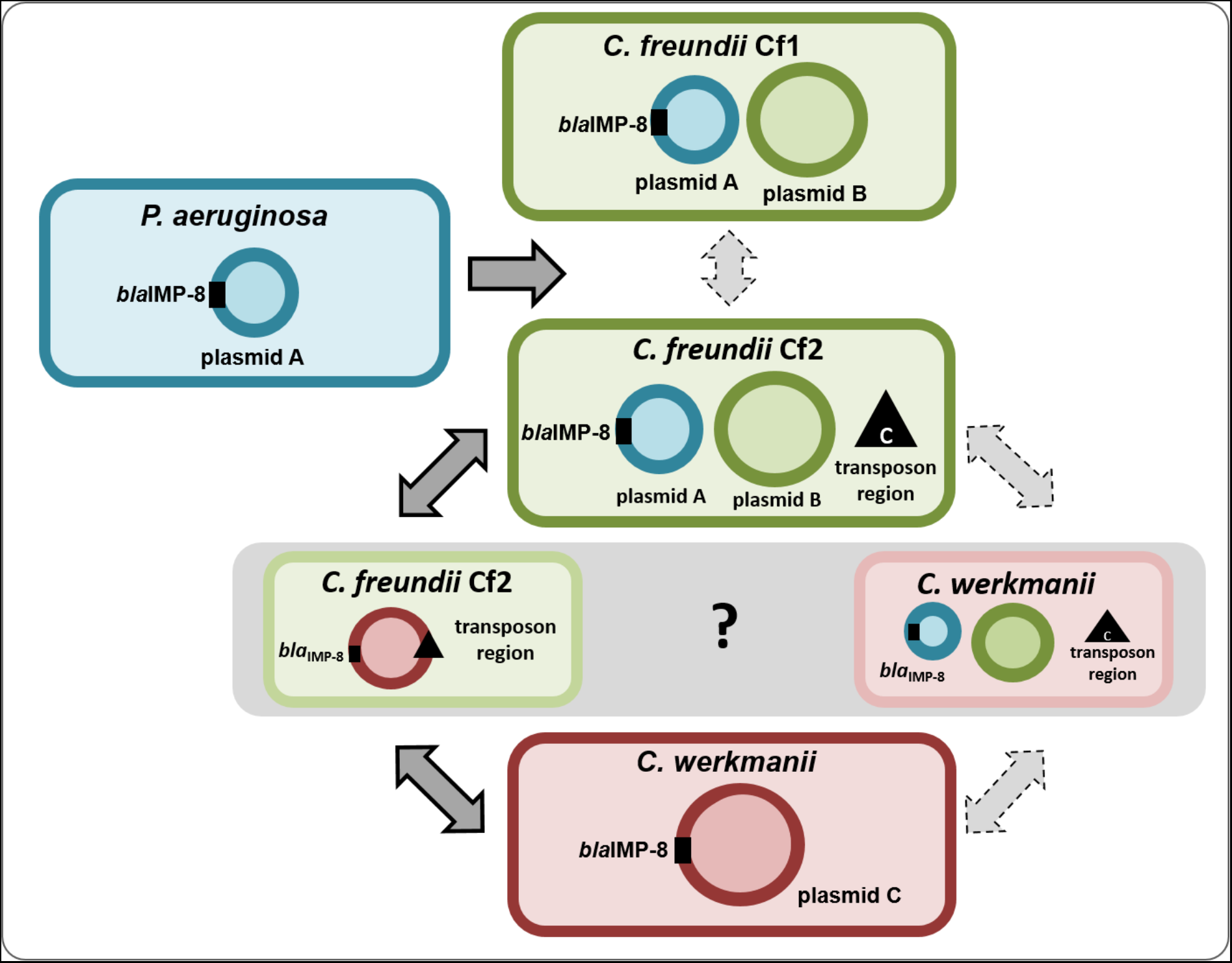
Concept of plasmid evolution and transmission across three bacterial species. *P. aeruginosa bla*_IMP-8_ was isolated approx. 2.5 years prior to the first isolation of *Citrobacter species* harboring *bla*_IMP-8_, leading to the hypothesis of a transfer of plasmid A from *P. aeruginosa* to *Citrobacter species*. The *C. freundii* and C. werkmanii *bla*_IMP-8_ started to occur at the same time, thus the timeline does not suggest a specific direction of the transfer. However, the existence of the transposable element region in the chromosome of *C. freundii* cluster Cf2 (marked with a black triangle) makes it the most likely host of the merge of plasmid A and B, which are linked by two copies of the transposable element region. Solid arrows represent the transmission sequence resulting from this hypothesis.

In conclusion, we demonstrated the successful application of Nanopore sequencing to track the transmission and rapid evolution of an antibiotic resistance plasmids within and between multiple bacterial species in a comprehensive and systematic collection of multi-drug resistant Gram-negative bacteria obtained from a large cohort of high-risk patients and corresponding environment samples.

## Discussion

Understanding the evolution and spread of multidrug resistant organisms has become a major challenge in the medical field, necessitating the development of novel diagnostic methods in order to effectively combat increasing numbers of infections with these organisms. The clinical importance of an antimicrobial resistance gene is determined by i) the class of antibiotics that is rendered resistant, ii) the pathogenicity of the bacterium and iii) the genetic location of the AMR gene. The localization of one or more AMR genes on a mobile genetic element, e.g. a plasmid, strongly increases the risk of resistance spreading between different bacterial genera, including well adapted and successful human pathogens established in the hospital environment.

In several studies, the spread of carbapenemase harboring plasmids has been demonstrated to happen in the hospital environment (e.g. [10]). For example, Conlan et al provided evidence for horizontal gene transfer of carbapenemase *bla*_KPC_ harboring plasmids between *Klebsiella pneumoniae*, *Enterobacter* and *Citrobacter* [10]. Interestingly, the *Citrobacter* strain from their study (CFNIH1), which was isolated from the hospital environment, contained a 272 kb plasmid harboring the carbapenemase gene KPC and clustered very closely with our study *C. werkmanii* isolate P28 (Figure 2C), which harbored the large plasmid C. This might indicate that the genetic background of this *Citrobacter* strains allows for large plasmid uptake or formation of megaplasmids in this species. The formation of multi-drug resistance megaplasmids has also been noted in other Enterobacteriaceae. For example, Desmet et al. analysed two clinical isolates harboring *bla*_OXA-427_ carbapenemase and identified a 321 kb megaplasmid which resulted from a cointegration of the MDR plasmid in another plasmid background [25]. A further study demonstrated, that a fusion plasmid had occurred as a result of recombination in a clinical Escherichia coli isolate containing the carbapenemase *bla*_NDM-5_ [26]. Of note, this *bla*_NDM-5_ megaplasmid harbored duplicate transposon regions in the plasmid, which resulted from the plasmid fusion. On the other hand, the fusion plasmid was not stable, when transferred to an *E. coli* recipient strain [26]. This is in line with our observation. Fusion plasmid C contained a duplicate transposon region, most likely as a result of recombination. While plasmid C was stable within our study isolates, it was never detected afterwards, suggesting that the plasmid was not positively selected in the hospital environment. However, further studies are needed to elucidate the factors involved in megaplasmid evolution dynamics.

Although the importance of plasmid evolution and horizontal gene transfer for the spread of MDR bacteria has clearly been documented, the epidemiological surveillance within hospitals is not commonly performed and remains limited to very few centers. Short-read sequencing technology, which is available in many hospitals, cannot reliably distinguish between plasmids and chromosomes and often leads to fragmented genome and plasmid assemblies. Long-read sequencing technologies on the other hand allow for high quality, finished assemblies of plasmids. With the emergence of Nanopore sequencing, a fast and inexpensive alternative technology for de novo assembly of multidrug resistant bacteria isolates became available [11, 13, 27]. Here, we demonstrated that the application of Nanopore sequencing in combination with Illumina short reads and epidemiological data allowed for detailed tracking of plasmid evolution on a comprehensive consecutive collection of *bla*_IMP-8_ harboring multi-drug resistant Gram-negative bacteria. In addition to multiple plasmid-based horizontal gene transfers, we were able to detect rearrangements within the multidrug resistance gene cassette, as well as fusion of two plasmids to a mega-plasmid. While the presence and absence of antimicrobial resistance genes can be postulated based on Illumina short read assemblies, their location on mobile elements and the determination of the structure of multidrug resistance gene cassettes remains challenging due to difficulties with assembling repetitive regions. In the *P. aeruginosa* genomes assembled using Nanopore data, we could readily detect continuous reads confirming the circularity of the plasmid and the exact order of the resistance gene cassette, and could distinguish between bacteria harboring the mega-plasmid or the two independent plasmids, further emphasizing the power of long reads for determination of structures of mobile genetic elements.

In the context of this study, we developed a Nanopore-based analysis pipeline for clinical isolates (termed *pathoLogic*) including the plasmid analysis method *plasmIDent*, in order to obtain richly annotated circularized plasmids and possible transmission, fusion or rearrangement events. Importantly, the pipeline is fully automated and required no manual curation of plasmid sequences. It supports pure Nanopore and Nanopore-Illumina hybrid assembly approaches, evaluates the circularity of plasmids, annotates resistance genes and performs a comparative genome and plasmid analysis between any number of hospital isolates suspected to harbor a specific resistance gene or resistance gene cassette. The chosen method for tracking of MDR resistance plasmids and their evolutionary dynamics represents a powerful approach, which could be applied for real-time infection control surveillance, in small scale using Nanopore MinION or in large-scale using Nanopore PromethION machines. The rapid detection of resistance spread via plasmids will allow for successful countermeasures and efficient containment of hospital outbreaks.

## Conclusion

The application of Nanopore sequencing and the establishment of a computational pipeline for genome and plasmid assembly, annotation and comparative analysis enabled us to investigate plasmid-driven adaptation and emergence of multidrug resistant bacteria using a comprehensive strain collection including patient and environment isolates. Using Nanopore-based de novo assemblies, we demonstrated that horizontal gene transfer via a multidrug resistance plasmid between *P. aeruginosa*, *C. freundii* and *C. werkmanii*, plasmid fusion resulting in a megaplasmid and evolution of the multidrug resistance gene cassette had occurred within the short period of 3 years within our hospital. In summary, we developed and showcased a novel pipeline for de novo bacterial genome assembly, AMR gene and plasmid characterisation and comparative analysis across species, enabling rapid tracking of AMR transmission via plasmids in hospital settings.

## Methods

### Study isolates

In total, 54 hospital strains were included in the study, comprising *P. aeruginosa* (n=45), *C. freundii* (n=8) and *C. werkmanii* (n=1). Strains were obtained from patient specimens including rectal screening culture (n=40) and water-related environment sources (toilet, sink, n=14). All isolates were cultured and identified following standard microbiology protocols as described before [28] and were positive for the *bla*_IMP-8_ gene as determined by PCR [29]. All isolates were recovered from samples of the haemato-oncology department between July 2009 and July 2015. During this time the sampling strategy for screening cultures and environmental surveillance was adjusted as a consequence of the *P. aeruginosa bla*_IMP-8_ outbreak. Between July 2009 and October 2010 only clinical specimens were obtained. Weekly rectal screening programs of all haemato-oncology patients and environment screening of toilets, sinks and showers in a 14-day turn were introduced in October 2010.

### Nanopore and Illumina sequencing

Nanopore sequencing was performed on Oxford Nanopore Technologies MinION device with three different chemistries (versions 6, 7 and 8) and flowcell versions (FLO-MAP103 version Pk.1, FLO-MIN105 version R9 and FLO-MIN106 version R9.4). An overview of chemistry and flowcell versions used for each sample is shown in supplementary table 2.

#### ONT chemistry version 6

Sequencing libraries were prepared with the Genomic DNA Sequencing SQK–MAP006 kit using 1.5 μg of gDNA as starting material. Briefly, the nick-repaired DNA (NEBNext FFPE DNA Repair Mix, NEB) was sheared in a Covaris g-TUBE (Covaris, Inc.), followed by end-repair and dA-tailing (NEBNext UltraII End Repair/dA-Tailing Module, NEB). The leader and hairpin sequencing adapters (ONT) were ligated using Blunt TA Ligase (NEB). After tether addition, the final library was purified with MyOne Streptavidin C1 Beads (Thermo Fisher). The MinION flowcell (FLO-MAP103, ONT) was primed and loaded with the library for 48h run with 24h interval of adding new pre-sequencing mix, running buffer and Fuel Mix (ONT).

#### ONT chemistry version 7 and 8

Libraries were prepared with the Genomic DNA Sequencing Kit SQK_NSK007 and SQK-LSK108 starting with 1.5 μg of gDNA sheared in a Covaris g-TUBE (Covaris, Inc.) and nick-repaired with NEBNext FFPE DNA Repair Mix (NEB). Subsequently, DNA was end-repaired and adenylated (NEBNext Ultra II End-Repair/dA-tailing Module, NEB) followed by ligation of adaptor (ONT) using NEB Blunt/TA Master Mix (NEB). After flowcells priming, FLO-MIN105 for kit SQK_NSK007 and FLO-MIN106 for kit SQK-LSK108 libraries were loaded and run for 48h following the manufacturer protocols (ONT).

#### Illumina sequencing

Due to the advances in sequencing technology available over the study period, different protocols were used to obtain short-read sequences, as described before [17, 28, 30]. In brief, early isolates were sequenced with 2x 50 bps on a Illumina HiSeq2000 [17], or by 2×300 bps on a Illumina Miseq [30] or using 2×250 bps on a Illumina Miseq [28]. Supplementary table 2 provides a detailed overview on the sequencing protocols applied.

### Hybrid *de novo* assembly pipeline using long and short reads

To achieve complete de novo genome assemblies we developed a custom pipeline (termed *pathoLogic*, see figure 1) consisting of individual steps for read pre-processing, hybrid de novo assembly, quality control and generation of assembly statistics. First, long Nanopore reads are subjected to adapter trimming with Porechop (https://github.com/rrwick/Porechop), quality filtering with Filtlong (https://github.com/rrwick/Filtlong) and quality control (QC) using Nanoplot [31]. Adapter trimming and QC for short reads is performed using SeqPurge [32]. We benchmarked multiple assembly approaches implemented in *pathologic*. *Unicycler*, a hybrid assembler using short and long read reads [16], produced the longest contigs at high and low read coverage and was therefore used in this study. Finally, assembly statistics are calculated and contigs shorter than 2000 bp are removed. Application specific parameters are documented in the published source code and configuration file. All tools are included in the provided Docker image (release v1.0) available on github (*plasmIDent*: https://github.com/imgag/plasmIDent, *pathoLogic*: https://github.com/imgag/pathoLogic).

### Phylogenetic analysis

Assembly of the short read Illumina data for all studied isolates was performed using Spades version 3.7.0 [33], followed by alignment using *ProgressiveMauve* (version 2.3.1) [34] with a locally collinear block size of 1000 bp. Phage content was removed using *Phast* [35]. For phylogeny calculation IQ tree version 1.6.3 was applied using the UFboot mode with 1000 bootstraps and the modelFinder [36–38]. For visualisation *Figtree* version v1.4.2 (http://tree.bio.ed.ac.uk/software/figtree/) was applied. For calculation of the *Citrobacter* maximum likelihood phylogeny, the 9 study isolates and 14 reference genomes were included (*Citrobacter amalonaticus* Y19 (CP011132_1), *Citrobacter braakii* GTA-CB01 (JRHK01000001.1), *C. braakii* GTA-CB04 (JRHL01000001.1), *Citrobacter farmeri* GTC 1319 (NZ_BBMX01000031.1), *C. freundii* CFNIH1 (NZ_CP007557_1), *Citrobacter rodentium* ICC168 (NC_013716_1), *Citrobacter sedlakii* NBRC 105722 (NZ_BBNB01000030.1), *C. werkmanii* NBRC 105721 (NZ_BBMW01000009.1), *Citrobacter youngae* ATCC 29220 (NZ_GG730308.1), *Citrobacter koseri ATCC* BAA-895 (CP000822.1), *C. freundii* ATCC 8090 (JMTA01000001.1), *C. rodentium* ATCC 51459 (JXUN01000001.1), *C. amalonaticus* L8A (JMQQ01000001.1) and *C. werkmanii* DSMZ17579 [30]. For *P. aeruginosa*, the 45 study isolates and one *P. aeruginosa* blaVIM outgroup strain were included in the analysis.

### Plasmid detection and annotation

For most isolates the assembly produced one or few large chromosomal scaffolds along with several shorter contigs between 10 kb and 200 kb in length. The latter could either stem from complete circular plasmids or from fragments of the chromosome or plasmids. We therefore developed the *plasmIDent* tool, which uses long reads to ascertain whether a scaffold is circular, identifies all antibiotic resistance genes and calculates characteristic metrices such as GC content and read coverage. *PlasmIDent* takes assembled genomes in fasta format and nanopore reads in fastq format as input. First, contig ends are fused in order to mimic a circular layout. Next, *minimap2* is used to align Nanopore reads to the putative plasmid and the end-end fusion site. If long reads continuously cover the scaffold and the artificially closed gap, we assume that the sequence originates from a circular plasmid. Furthermore, sudden changes of median GC content within the plasmid are used to predict ancestral fusions of two or multiple plasmids. Finally, *plasmIDent* supports discovery of resistance genes using the CARD database.

### Genome annotations

Assembled FASTA files were uploaded to the ResFinder tool (https://cge.cbs.dtu.dk//services/ResFinder/) applying a 98 % identity threshold and a minimum overlapping length of 60%. Additionally, CARD-based annotations automatically generated by *plasmIDent* were merged with the *ResFinder* results. Finally, we used the RAST Webservice to get complete genome and plasmid annotations for all isolates and the *ISFinder* Webservice to specifically identify transposable elements.

### Comparative genome and plasmid analysis across species

#### Whole genome alignment (WGA)

Multiple whole genome alignments of all assembled plasmids were generated with *progressiveMauve* in order to find highly similar regions. Plasmids with highly homologous regions were additionally compared by pairwise sequence alignment using nucmer (e. g. Fig S3), resulting in a pairwise identity score and the annotation of homologous regions. We used dotplots (*pathoLogic* utility scripts) of the pairwise alignments to visually identify rearrangements in plasmids. Homologous regions between plasmids and chromosomal scaffolds were identified using pairwise alignment (nucmer) between a plasmid of interest against the concatenated sequence of all scaffolds in an isolate’s genome assembly. More specifically, we identified homologous sequences of the transposable element region found in plasmid C, but not A and B in order to ascertain, if a Citrobacter isolate contains only plasmids A and B and the transposon region inserted in the chromosome, or if it contains plasmid C with the transposon region in the plasmid.

#### Read coverage (density) analysis

We chose megaplasmid C of isolate 28_P_CW as reference plasmid, as it integrates both plasmids A and B involved in the studied horizontal gene transfer of AMR genes. We used bwa-mem to realign Illumina short reads of each isolate to the reference plasmid, thereby identifying the presence or absence of specific regions based on read density (i.e. regions without read coverage are absent in a studied isolate, see figures 4 and S4). We identified breakpoints, indicating structural variants or the end of plasmids, based on clip- or split-reads (see figure S2). We defined deletions as regions with very low density read coverage, with split- or paired-reads spanning the two breakpoints. (Plasmid ends are identified by circularization as described before.) Furthermore, we evaluated that putatively deleted resistance genes were also absent from the plasmid AMR gene annotations by ResFinder and CARD.

#### AMR gene rearrangements

WGA of the resistance gene cassette of all isolates assembled with Nanopore reads identified two haplotypes termed RSC1 and RSC2, distinguished by two translocations of AMR genes. In order to assign all sequenced isolates to one or the other cassette configuration we aligned Illumina short reads to the 4 breakpoints per haplotype (two breakpoints for each translocation event per cassette configuration). Then, we compared the number of aligned reads spanning the four breakpoints in RSC1 vs. RSC2 and computed the log transformed fraction of breaks in RSC1 and RSC2, each normalized by the respective amount of total reads. Isolates showing log-values > 1 were assigned to RSC1 and log-values < −1 to RSC2, while other isolates remained unassigned.

## Declarations

### Ethics approval and consent to participate

The study was conducted in accordance with the local ethic committee (Ethic committee Medical faculty, University of Tübingen, No. 741/2016BO2 and 407/2013R).

### Consent for publication

Not applicable.

### Availability of data and material

All sequence data is deposited at the European Nucleotide Archive (accession number PRJEB31907, https://www.ebi.ac.uk/ena/data/view/PRJEB31907). *PathoLogic* (https://github.com/imgag/pathoLogic) and *plasmIDent* (https://github.com/imgag/plasmIDent) developed in this study are freely available on Github.

### Authors’ contributions

SP, DB, PO, JG, MM, MW, IG, MG generated the laboratory and sequencing data. JL, WV and DD gathered epidemiological data. SP, MB, CG and SO performed the data analysis. CG, MB and SO developed the bioinformatics methods and pipelines, LB developed the GUI. SP, IA and SO designed the study. SP, CG and SO wrote the manuscript.

## Acknowledgements

We thank Nadine Hoffmann and Baris Bader for expert technical assistance.

## References

1. Roca I, Akova M, Baquero F, Carlet J, Cavaleri M, Coenen S, Cohen J, Findlay D, Gyssens I, Heuer OE, et al: The global threat of antimicrobial resistance: science for intervention. New Microbes New Infect 2015, 6:22–29.

2. Maraolo AE, Cascella M, Corcione S, Cuomo A, Nappa S, Borgia G, De Rosa FG, Gentile I: Management of multidrug-resistant Pseudomonas aeruginosa in the intensive care unit: state of the art. Expert Rev Anti Infect Ther 2017, 15:861–871.

3. Kengkla K, Kongpakwattana K, Saokaew S, Apisarnthanarak A, Chaiyakunapruk N: Comparative efficacy and safety of treatment options for MDR and XDR Acinetobacter baumannii infections: a systematic review and network meta-analysis. J Antimicrob Chemother 2018, 73:22–32.

4. Tacconelli E, Cataldo MA, Dancer SJ, De Angelis G, Falcone M, Frank U, Kahlmeter G, Pan A, Petrosillo N, Rodriguez-Bano J, et al: ESCMID guidelines for the management of the infection control measures to reduce transmission of multidrug-resistant Gram-negative bacteria in hospitalized patients. Clin Microbiol Infect 2014, 20 Suppl 1:1–55.

5. Naylor NR, Atun R, Zhu N, Kulasabanathan K, Silva S, Chatterjee A, Knight GM, Robotham JV: Estimating the burden of antimicrobial resistance: a systematic literature review. Antimicrob Resist Infect Control 2018, 7:58.

6. Gilchrist CA, Cotton JA, Burkey C, Arju T, Gilmartin A, Lin Y, Ahmed E, Steiner K, Alam M, Ahmed S, et al: Genetic Diversity of Cryptosporidium hominis in a Bangladeshi Community as Revealed by Whole-Genome Sequencing. J Infect Dis 2018, 218:259–264.

7. Partridge SR, Kwong SM, Firth N, Jensen SO: Mobile Genetic Elements Associated with Antimicrobial Resistance. Clin Microbiol Rev 2018, 31.

8. Arredondo-Alonso S, Willems RJ, van Schaik W, Schurch AC: On the (im)possibility of reconstructing plasmids from whole-genome short-read sequencing data. Microb Genom 2017, 3:e000128.

9. Conlan S, Park M, Deming C, Thomas PJ, Young AC, Coleman H, Sison C, Program NCS, Weingarten RA, Lau AF, et al: Plasmid Dynamics in KPC-Positive Klebsiella pneumoniae during Long-Term Patient Colonization. MBio 2016, 7.

10. Conlan S, Thomas PJ, Deming C, Park M, Lau AF, Dekker JP, Snitkin ES, Clark TA, Luong K, Song Y, et al: Single-molecule sequencing to track plasmid diversity of hospital-associated carbapenemase-producing Enterobacteriaceae. Sci Transl Med 2014, 6:254ra126.

11. Lemon JK, Khil PP, Frank KM, Dekker JP: Rapid Nanopore Sequencing of Plasmids and Resistance Gene Detection in Clinical Isolates. J Clin Microbiol 2017, 55:3530–3543.

12. George S, Pankhurst L, Hubbard A, Votintseva A, Stoesser N, Sheppard AE, Mathers A, Norris R, Navickaite I, Eaton C, et al: Resolving plasmid structures in Enterobacteriaceae using the MinION nanopore sequencer: assessment of MinION and MinION/Illumina hybrid data assembly approaches. Microb Genom 2017, 3:e000118.

13. Dong N, Yang X, Zhang R, Chan EW, Chen S: Tracking microevolution events among ST11 carbapenemase-producing hypervirulent Klebsiella pneumoniae outbreak strains. Emerg Microbes Infect 2018, 7:146.

14. Chin CS, Alexander DH, Marks P, Klammer AA, Drake J, Heiner C, Clum A, Copeland A, Huddleston J, Eichler EE, et al: Nonhybrid, finished microbial genome assemblies from long-read SMRT sequencing data. Nat Methods 2013, 10:563–569.

15. Antipov D, Korobeynikov A, McLean JS, Pevzner PA: hybridSPAdes: an algorithm for hybrid assembly of short and long reads. Bioinformatics 2016, 32:1009–1015.

16. Wick RR, Judd LM, Gorrie CL, Holt KE: Unicycler: Resolving bacterial genome assemblies from short and long sequencing reads. PLoS Comput Biol 2017, 13:e1005595.

17. Willmann M, Bezdan D, Zapata L, Susak H, Vogel W, Schroppel K, Liese J, Weidenmaier C, Autenrieth IB, Ossowski S, Peter S: Analysis of a long-term outbreak of XDR Pseudomonas aeruginosa: a molecular epidemiological study. J Antimicrob Chemother 2015, 70:1322–1330.

18. Peter S, Wolz C, Kaase M, Marschal M, Schulte B, Vogel W, Autenrieth I, Willmann M: Emergence of Citrobacter freundii carrying IMP-8 metallo-beta-lactamase in Germany. New Microbes New Infect 2014, 2:42–45.

19. Di Tommaso P, Chatzou M, Floden EW, Barja PP, Palumbo E, Notredame C: Nextflow enables reproducible computational workflows. Nat Biotechnol 2017, 35:316–319.

20. Koren S, Walenz BP, Berlin K, Miller JR, Bergman NH, Phillippy AM: Canu: scalable and accurate long-read assembly via adaptive k-mer weighting and repeat separation. Genome Res 2017, 27:722–736.

21. Li H: Minimap and miniasm: fast mapping and de novo assembly for noisy long sequences. Bioinformatics 2016, 32:2103–2110.

22. Kolmogorov M, Yuan J, Lin Y, Pevzner PA: Assembly of Long Error-Prone Reads Using Repeat Graphs. bioRxiv 2018.

23. Lin Y, Yuan J, Kolmogorov M, Shen MW, Chaisson M, Pevzner PA: Assembly of long error-prone reads using de Bruijn graphs. Proc Natl Acad Sci U S A 2016, 113:E8396–E8405.

24. Boyd ES, Barkay T: The mercury resistance operon: from an origin in a geothermal environment to an efficient detoxification machine. Front Microbiol 2012, 3:349.

25. Desmet S, Nepal S, van Dijl JM, Van Ranst M, Chlebowicz MA, Rossen JW, Van Houdt JKJ, Maes P, Lagrou K, Bathoorn E: Antibiotic Resistance Plasmids Cointegrated into a Megaplasmid Harboring the blaOXA-427 Carbapenemase Gene. Antimicrob Agents Chemother 2018, 62.

26. Xie M, Li R, Liu Z, Chan EWC, Chen S: Recombination of plasmids in a carbapenem-resistant NDM-5-producing clinical Escherichia coli isolate. J Antimicrob Chemother 2018, 73:1230–1234.

27. Li R, Xie M, Dong N, Lin D, Yang X, Wong MHY, Chan EW, Chen S: Efficient generation of complete sequences of MDR-encoding plasmids by rapid assembly of MinION barcoding sequencing data. Gigascience 2018, 7:1–9.

28. Peter S, Oberhettinger P, Schuele L, Dinkelacker A, Vogel W, Dorfel D, Bezdan D, Ossowski S, Marschal M, Liese J, Willmann M: Genomic characterisation of clinical and environmental Pseudomonas putida group strains and determination of their role in the transfer of antimicrobial resistance genes to Pseudomonas aeruginosa. BMC Genomics 2017, 18:859.

29. Pitout JD, Gregson DB, Poirel L, McClure JA, Le P, Church DL: Detection of Pseudomonas aeruginosa producing metallo-beta-lactamases in a large centralized laboratory. J Clin Microbiol 2005, 43:3129–3135.

30. Peter S, Bezdan D, Oberhettinger P, Vogel W, Dorfel D, Dick J, Marschal M, Liese J, Weidenmaier C, Autenrieth I, et al: Whole-genome sequencing enabling the detection of a colistin-resistant hypermutating Citrobacter werkmanii strain harbouring a novel metallo-beta-lactamase VIM-48. Int J Antimicrob Agents 2018, 51:867–874.

31. De Coster W, D’Hert S, Schultz DT, Cruts M, Van Broeckhoven C: NanoPack: visualizing and processing long-read sequencing data. Bioinformatics 2018, 34:2666–2669.

32. Sturm M, Schroeder C, Bauer P: SeqPurge: highly-sensitive adapter trimming for paired-end NGS data. BMC Bioinformatics 2016, 17:208.

33. Bankevich A, Nurk S, Antipov D, Gurevich AA, Dvorkin M, Kulikov AS, Lesin VM, Nikolenko SI, Pham S, Prjibelski AD, et al: SPAdes: a new genome assembly algorithm and its applications to single-cell sequencing. J Comput Biol 2012, 19:455–477.

34. Darling AE, Mau B, Perna NT: progressiveMauve: multiple genome alignment with gene gain, loss and rearrangement. PLoS One 2010, 5:e11147.

35. Zhou Y, Liang Y, Lynch KH, Dennis JJ, Wishart DS: PHAST: a fast phage search tool. Nucleic Acids Res 2011, 39:W347–352.

36. Nguyen LT, Schmidt HA, von Haeseler A, Minh BQ: IQ-TREE: a fast and effective stochastic algorithm for estimating maximum-likelihood phylogenies. Mol Biol Evol 2015, 32:268–274.

37. Hoang DT, Chernomor O, von Haeseler A, Minh BQ, Vinh LS: UFBoot2: Improving the Ultrafast Bootstrap Approximation. Mol Biol Evol 2018, 35:518–522.

38. Kalyaanamoorthy S, Minh BQ, Wong TKF, von Haeseler A, Jermiin LS: ModelFinder: fast model selection for accurate phylogenetic estimates. Nat Methods 2017, 14:587–589.

