## Supplementary Materials and Methods for "Tracking of antibiotic resistance transfer and rapid plasmid evolution in a hospital setting by Nanopore sequencing"

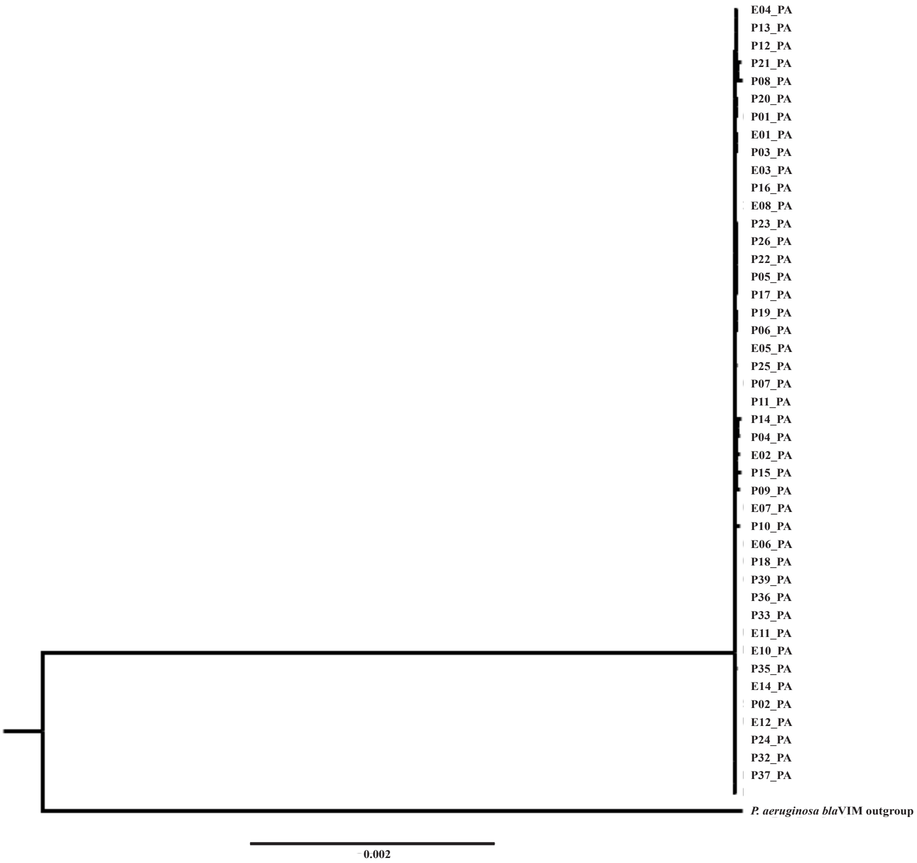


**Figure S1.** Maximum likelihood phylogeny of *P. aeruginosa species* (n=45) included in the study. All study isolates are closely clustered. A *P. aeruginosa bla*$_{VIM}$ strain was included as outgroup. The scale bar shows the expected number of nucleotide changes per site.

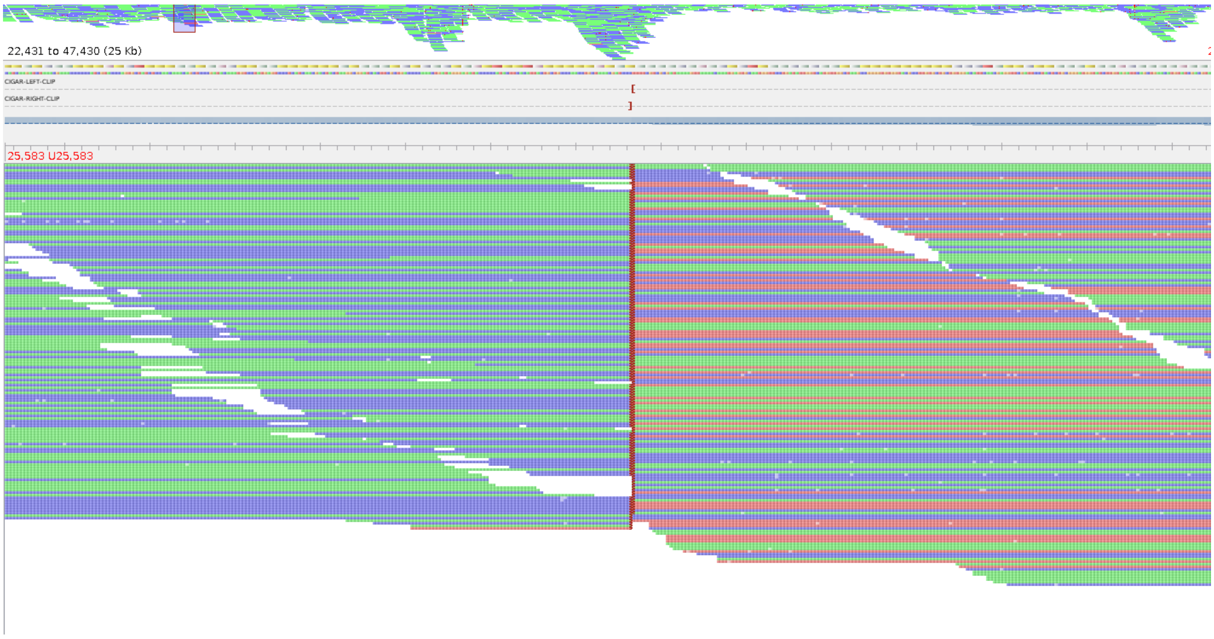


**Figure S2.** Visualisation of Illumina reads from *C. freundii* isolates mapped to the reference plasmid C of *C. werkmanii* using the *Tablet* software [1]. Despite no obvious changes in coverage, a distinct break is visible where all reads are clipped. Blue and green colour shows the direction of the aligned read. Reads coloured in red indicate that the mate of the read pair is not mapped. These manually curated breaks are included in figure 4A as red lines.

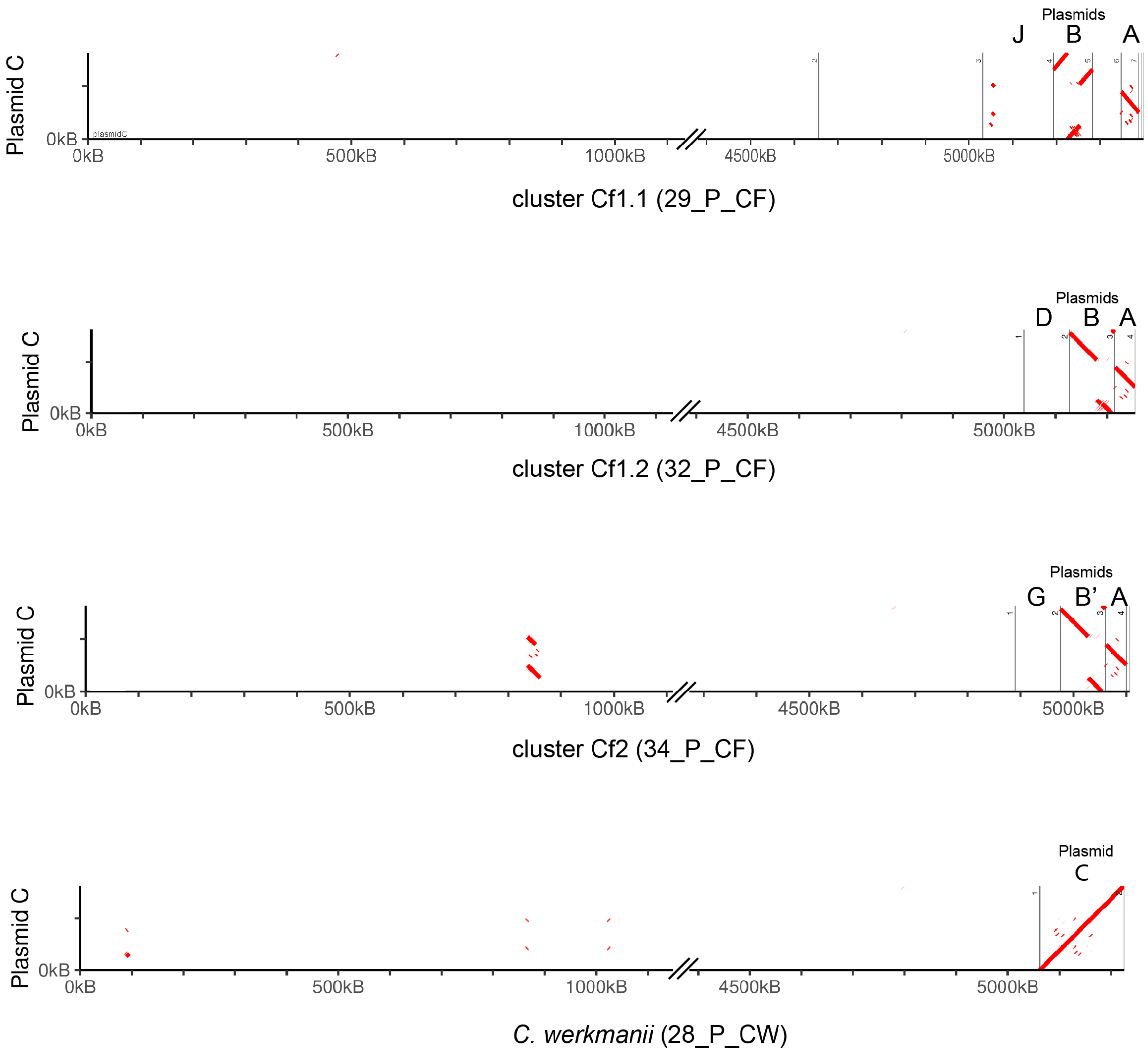


**Figure S3.** Dotplot visualisation of plasmid C (y-axis) aligned to the complete genomes and plasmids of four representative isolates (x-axis). Regions without alignments are hidden. The alignment of *C. freundii* isolate 34_P_CF (Cf2) shows that parts of plasmid C match to a region on the chromosome at ca. 850 kbp. This indicates that the Tn3 regions fusing plasmid A and B to C likely originated in the chromosomes of *Citrobacter freundii* cluster Cf2.

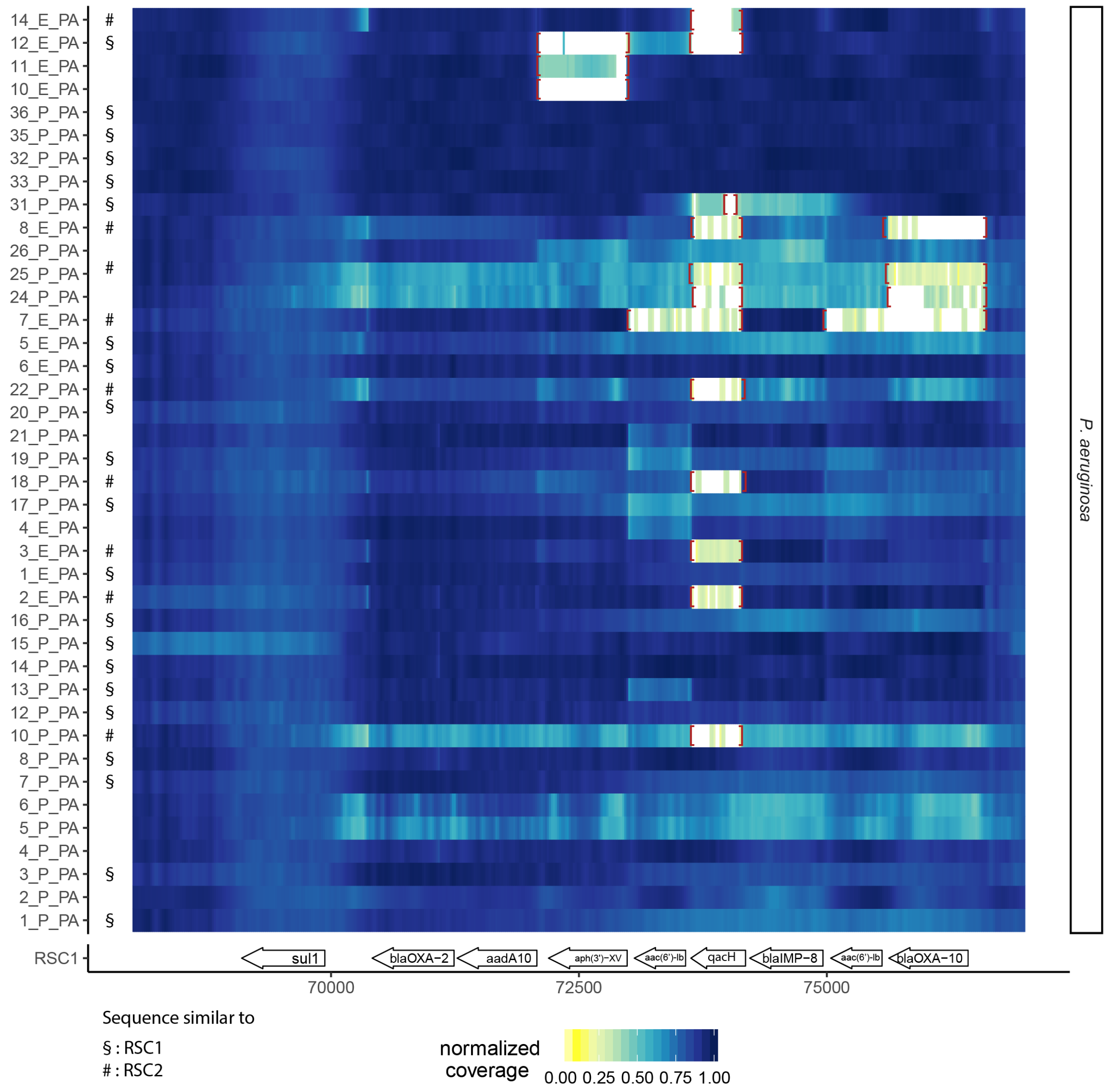


**Figure S4.** Coverage plot of short read Illumina data mapped against plasmid C. Presented here are only *P. aeruginosa* strains for which no nanopore sequencing data was available. The selected region focuses on the resistance cassette (reference is cassette configuration RSC1), highlighting a high number of deletions and coverage variations between different samples. Manually confirmed borders of deleted regions (as showcased in figure S2) are included as red brackets (see also Figure 4 for coverage plots of samples for which Nanopore data was available).

| Strain_ID | Bacterial species | Library insert size | Read Length | Sequencing Platform | Read 1 File | Read 2 File |
|---|---|---|---|---|---|---|
| 1_P_PA | *P . aeruginosa* | 300 | 2x50 | HiSeq2000 | 1_P_PA_1.fastq.gz | 1_P_PA_2.fastq.gz |
| 2_P_PA | *P . aeruginosa* | 800 | 2x300 | MiSeq | 2_P_PA_1.fastq.gz | 2_P_PA_2.fastq.gz |
| 3_P_PA | *P . aeruginosa* | 300 | 2x50 | HiSeq2000 | 3_P_PA_1.fastq.gz | 3_P_PA_2.fastq.gz |
| 4_P_PA | *P . aeruginosa* | 300 | 2x50 | HiSeq2000 | 4_P_PA_1.fastq.gz | 4_P_PA_2.fastq.gz |
| 5_P_PA | *P . aeruginosa* | 300 | 2x50 | HiSeq2000 | 5_P_PA_1.fastq.gz | 5_P_PA_2.fastq.gz |
| 6_P_PA | *P . aeruginosa* | 300 | 2x50 | HiSeq2000 | 6_P_PA_1.fastq.gz | 6_P_PA_2.fastq.gz |
| 7_P_PA | *P . aeruginosa* | 300 | 2x50 | HiSeq2000 | 7_P_PA_1.fastq.gz | 7_P_PA_2.fastq.gz |
| 8_P_PA | *P . aeruginosa* | 300 | 2x50 | HiSeq2000 | 8_P_PA_1.fastq.gz | 8_P_PA_2.fastq.gz |
| 9_P_PA | *P . aeruginosa* | 300 | 2x50 | HiSeq2000 | 9_P_PA_1.fastq.gz | 9_P_PA_2.fastq.gz |
| 9_P_PA | *P . aeruginosa* | | | Nanopore | 9_P_PA_nano.fastq.gz | n.a. |
| 10_P_PA | *P . aeruginosa* | 300 | 2x50 | HiSeq2000 | 10_P_PA_1.fastq.gz | 10_P_PA_2.fastq.gz |
| 11_P_PA | *P . aeruginosa* | 300 | 2x50 | HiSeq2000 | 11_P_PA_1.fastq.gz | 11_P_PA_2.fastq.gz |
| 11_P_PA | *P . aeruginosa* | | | Nanopore | 11_P_PA_nano.fastq.gz | |
| 12_P_PA | *P . aeruginosa* | 300 | 2x50 | HiSeq2000 | 12_P_PA_1.fastq.gz | 12_P_PA_2.fastq.gz |
| 13_P_PA | *P . aeruginosa* | 300 | 2x50 | HiSeq2000 | 13_P_PA_1.fastq.gz | 13_P_PA_2.fastq.gz |
| 14_P_PA | *P . aeruginosa* | 300 | 2x50 | HiSeq2000 | 14_P_PA_1.fastq.gz | 14_P_PA_2.fastq.gz |
| 15_P_PA | *P . aeruginosa* | 300 | 2x50 | HiSeq2000 | 15_P_PA_1.fastq.gz | 15_P_PA_2.fastq.gz |
| 16_P_PA | *P . aeruginosa* | 300 | 2x50 | HiSeq2000 | 16_P_PA_1.fastq.gz | 16_P_PA_2.fastq.gz |
| 17_P_PA | *P . aeruginosa* | 300 | 2x50 | HiSeq2000 | 17_P_PA_1.fastq.gz | 17_P_PA_2.fastq.gz |
| 18_P_PA | *P . aeruginosa* | 300 | 2x50 | HiSeq2000 | 18_P_PA_1.fastq.gz | 18_P_PA_2.fastq.gz |
| 19_P_PA | *P . aeruginosa* | 300 | 2x50 | HiSeq2000 | 19_P_PA_1.fastq.gz | 19_P_PA_2.fastq.gz |
| 20_P_PA | *P . aeruginosa* | 300 | 2x50 | HiSeq2000 | 20_P_PA_1.fastq.gz | 20_P_PA_2.fastq.gz |
| 21_P_PA | *P . aeruginosa* | 300 | 2x50 | HiSeq2000 | 21_P_PA_1.fastq.gz | 21_P_PA_2.fastq.gz |
| 22_P_PA | *P . aeruginosa* | 300 | 2x50 | HiSeq2000 | 22_P_PA_1.fastq.gz | 22_P_PA_2.fastq.gz |
| 23_P_PA | *P . aeruginosa* | 300 | 2x50 | HiSeq2000 | 23_P_PA_1.fastq.gz | 23_P_PA_2.fastq.gz |
| 23_P_PA | *P . aeruginosa* | | | Nanopore | 23_P_PA_nano.fastq.gz | |
| 24_P_PA | *P . aeruginosa* | 300 | 2x50 | HiSeq2000 | 24_P_PA_1.fastq.gz | 24_P_PA_2.fastq.gz |
| 25_P_PA | *P . aeruginosa* | 300 | 2x50 | HiSeq2000 | 25_P_PA_1.fastq.gz | 25_P_PA_2.fastq.gz |
| 26_P_PA | *P . aeruginosa* | 300 | 2x50 | HiSeq2000 | 26_P_PA_1.fastq.gz | 26_P_PA_2.fastq.gz |
| 27_P_CF | *C. freundii* | 550 | 2x250 | MiSeq | 27_P_CF_1.fastq.gz | 27_P_CF_2.fastq.gz |
| 27_P_CF | *C. freundii* | | | Nanopore | 27_P_CF_nano.fastq.gz | n.a. |
| 28_P_CW | *C. werkmanii* | 800 | 2x300 | MiSeq | 28_P_CW_1.fastq.gz | 28_P_CW_2.fastq.gz |
| 28_P_CW | *C. werkmanii* | | | Nanopore | 28_P_CW_nano.fastq.gz | n.a. |
| 29_P_CF | *C. freundii* | 550 | 2x250 | MiSeq | 29_P_CF_1.fastq.gz | 29_P_CF_2.fastq.gz |
| 29_P_CF | *C. freundii* | | | Nanopore | 29_P_CF_nano.fastq.gz | n.a. |
| 30_P_CF | *C. freundii* | 550 | 2x250 | MiSeq | 30_P_CF_1.fastq.gz | 30_P_CF_2.fastq.gz |
| 30_P_CF | *C. freundii* | | | Nanopore | 30_P_CF_nano.fastq.gz | n.a. |
| 31_P_PA | *P . aeruginosa* | 550 | 2x250 | MiSeq | 31_P_PA_1.fastq.gz | 31_P_PA_2.fastq.gz |
| 32_P_CF | *C. freundii* | 550 | 2x250 | MiSeq | 32_P_CF_1.fastq.gz | 32_P_CF_2.fastq.gz |
| 32_P_CF | *C. freundii* | | | Nanopore | 32_P_CF_nano.fastq.gz | n.a. |

| | | | | | | |
|---|---|---|---|---|---|---|
| 32_P_PA | *P . aeruginosa* | 550 | 2x250 | MiSeq | 32_P_PA_1.fastq.gz | 32_P_PA_2.fastq.gz |
| 33_P_PA | *P . aeruginosa* | 550 | 2x250 | MiSeq | 33_P_PA_1.fastq.gz | 33_P_PA_2.fastq.gz |
| 34_P_CF | *C. freundii* | 550 | 2x250 | MiSeq | 34_P_CF_1.fastq.gz | 34_P_CF_2.fastq.gz |
| 34_P_CF | *C. freundii* | | | Nanopore | 34_P_CF_nano.fastq.gz | n.a. |
| 35_P_PA | *P . aeruginosa* | 550 | 2x250 | MiSeq | 35_P_PA_1.fastq.gz | 35_P_PA_2.fastq.gz |
| 36_P_PA | *P . aeruginosa* | 550 | 2x250 | MiSeq | 36_P_PA_1.fastq.gz | 36_P_PA_2.fastq.gz |
| 37_P_PA | *P . aeruginosa* | 550 | 2x250 | MiSeq | 37_P_PA_1.fastq.gz | 37_P_PA_2.fastq.gz |
| 37_P_PA | *P . aeruginosa* | | | Nanopore | 37_P_PA_nano.fastq.gz | n.a. |
| 38_P_CF | *C. freundii* | 550 | 2x250 | MiSeq | 38_P_CF_1.fastq.gz | 38_P_CF_2.fastq.gz |
| 38_P_CF | *C. freundii* | | | Nanopore | 38_P_CF_nano.fastq.gz | n.a. |
| 39_P_PA | *P . aeruginosa* | 550 | 2x250 | MiSeq | 39_P_PA_1.fastq.gz | 39_P_PA_2.fastq.gz |
| 39_P_PA | *P . aeruginosa* | | | Nanopore | 39_P_PA_nano.fastq.gz | n.a. |
| 1_E_PA | *P . aeruginosa* | 300 | 2x50 | HiSeq2000 | 1_E_PA_1.fastq.gz | 1_E_PA_2.fastq.gz |
| 2_E_PA | *P . aeruginosa* | 300 | 2x50 | HiSeq2000 | 2_E_PA_1.fastq.gz | 2_E_PA_2.fastq.gz |
| 3_E_PA | *P . aeruginosa* | 300 | 2x50 | HiSeq2000 | 3_E_PA_1.fastq.gz | 3_E_PA_2.fastq.gz |
| 4_E_PA | *P . aeruginosa* | 300 | 2x50 | HiSeq2000 | 4_E_PA_1.fastq.gz | 4_E_PA_2.fastq.gz |
| 5_E_PA | *P . aeruginosa* | 300 | 2x50 | HiSeq2000 | 5_E_PA_1.fastq.gz | 5_E_PA_2.fastq.gz |
| 6_E_PA | *P . aeruginosa* | 300 | 2x50 | HiSeq2000 | 6_E_PA_1.fastq.gz | 6_E_PA_2.fastq.gz |
| 7_E_PA | *P . aeruginosa* | 300 | 2x50 | HiSeq2000 | 7_E_PA_1.fastq.gz | 7_E_PA_2.fastq.gz |
| 8_E_PA | *P . aeruginosa* | 300 | 2x50 | HiSeq2000 | 8_E_PA_1.fastq.gz | 8_E_PA_2.fastq.gz |
| 9_E_CF | *C. freundii* | 550 | 2x250 | MiSeq | 9_E_CF_1.fastq.gz | 9_E_CF_2.fastq.gz |
| 9_E_CF | *C. freundii* | | | Nanopore | 9_E_CF_nano.fastq.gz | n.a. |
| 10_E_PA | *P . aeruginosa* | 550 | 2x250 | MiSeq | 10_E_PA_1.fastq.gz | 10_E_PA_2.fastq.gz |
| 11_E_PA | *P . aeruginosa* | 550 | 2x250 | MiSeq | 11_E_PA_1.fastq.gz | 11_E_PA_2.fastq.gz |
| 12_E_PA | *P . aeruginosa* | 550 | 2x250 | MiSeq | 12_E_PA_1.fastq.gz | 12_E_PA_2.fastq.gz |
| 13_E_CF | *C. freundii* | 550 | 2x250 | MiSeq | 13_E_CF_1.fastq.gz | 13_E_CF_2.fastq.gz |
| 13_E_CF | *C. freundii* | | | Nanopore | 13_E_CF_nano.fastq.gz | n.a. |
| 14_E_PA | *P . aeruginosa* | 550 | 2x250 | MiSeq | 14_E_PA_1.fastq.gz | 14_E_PA_2.fastq.gz |

**Table S1.** Overview of bacterial isolates and sequencing parameters. Isolate identifier (ID) (first column) encode the sample ID (sorted by sampling date), sampling source (E = environment, P = patient) and the species (PA = *P. aeruginosa*, CF = *C. freundii*, CW – *C. werkmanii*). Three different Illumina protocols and/or sequencing machines were applied over the course of six years, including HiSeq2000 with 2x 50 bps single-end reads [2], MiSeq with 2x300 bps [3], and MiSeq 2x250 bps paired-end reads [4]. Detailed descriptions of the sequencing procedures can be found in the corresponding references. Details on the applied ONT MinION sequencing protocols can be found in table S2 and Material and Methods.

| Sample | Number of reads: | Median read length: | Median read qual. | Flowcell version: | ONT Kit: |
|---|---|---|---|---|---|
| 28_P_CW | 107501 | 3731 | 7.7 | FLO-MAP103 | SQK_MAP006 |
| 30_P_CF | 449375 | 3300 | 9.4 | FLO-MIN106 | SQK-LSK108 |
| 32_P_CF | 689747 | 3134 | 9 | FLO-MIN106 | SQK-LSK108 |
| 34_P_CF | 640898 | 3163 | 9.8 | FLO-MIN106 | SQK-LSK108 |
| 9_E_CF | 59275 | 1572 | 7.8 | FLO-MIN106 | SQK-LSK108 |
| 38_P_CF | 327411 | 5908 | 8.6 | FLO-MIN106 | SQK-LSK108 |
| 27_P_CF | 89853 | 1918 | 6.9 | FLO-MIN105 | SQK-NSK007 |
| 29_P_CF | 926772 | 2753 | 9.7 | FLO-MIN106 | SQK-LSK108 |
| 13_E_CF | 224377 | 2136 | 9.1 | FLO-MIN106 | SQK-LSK108 |
| 9_P_PA | 71923 | 4373 | 8.5 | FLO-MIN105 | SQK-NSK007 |
| 11_P_PA | 50986 | 6222 | 8.8 | FLO-MIN105 | SQK-NSK007 |
| 37_P_PA | 137180 | 3542 | 6.5 | FLO-MIN105 | SQK-NSK007 |
| 39_P_PA | 41934 | 1340 | 3.9 | FLO-MIN105 | SQK-NSK007 |
| 23_P_PA | 149047 | 2127 | 6.6 | FLO-MIN106 | SQK-LSK108 |

**Table S2.** Quality parameters for ONT data obtained in this study, calculated using *Nanoplot* after adapter trimming with *Porechop* (https://github.com/rrwick/Porechop). Also shown are the Flowcell and Reagent Kit version used for the nanopore-sequencing run.

| Sample ID | # contigs | Largest contig | Total length | GC (%) | N50 | N75 | L50 | L75 |
|---|---|---|---|---|---|---|---|---|
| 28_P_CW | 3 | 5062321 | 5230854 | 52.09 | 5062321 | 5062321 | 1 | 1 |
| 30_P_CF | 6 | 4745085 | 5272837 | 51.88 | 4745085 | 4745085 | 1 | 1 |
| 32_P_CF | 5 | 5038088 | 5260813 | 51.87 | 5038088 | 5038088 | 1 | 1 |
| 34_P_CF | 5 | 4892588 | 5110715 | 51.71 | 4892588 | 4892588 | 1 | 1 |
| 9_E_CF | 29 | 1444960 | 5262884 | 51.80 | 636697 | 296195 | 3 | 6 |
| 38_P_CF | 6 | 4889818 | 5112060 | 51.69 | 4889818 | 4889818 | 1 | 1 |
| 27_P_CF | 25 | 1834568 | 5388324 | 51.84 | 915012 | 298121 | 2 | 5 |
| 29_P_CF | 10 | 2415109 | 5401457 | 51.80 | 2241041 | 2241041 | 2 | 2 |
| 13_E_CF | 11 | 3937886 | 5105413 | 51.69 | 3937886 | 3937886 | 1 | 1 |
| 9_P_PA | 18 | 1941814 | 6786480 | 66.14 | 1894580 | 572642 | 2 | 4 |
| 11_P_PA | 14 | 1894702 | 6808107 | 66.12 | 1693847 | 948850 | 2 | 4 |
| 37_P_PA | 20 | 1907170 | 6818725 | 66.11 | 1047546 | 547243 | 3 | 5 |
| 39_P_PA | 19 | 1988153 | 6650226 | 66.21 | 1549117 | 1486086 | 2 | 3 |
| 23_P_PA | 36 | 1047588 | 6766190 | 66.14 | 466918 | 315712 | 5 | 9 |

**Table S3.** Assembly quality parameters calculated with *QUAST* [5].

**A)**

| Strain Id | Species | # Plasmids | Plasmid types |
|---|---|---|---|
| 9_P_PA | *P. aeruginosa* | 4 | A , E, F |
| 11_P_PA | *P. aeruginosa* | 3 | A, E |
| 37_P_PA | *P. aeruginosa* | 3 | A, E |
| 39_P_PA | *P. aeruginosa* | 3 | A', E |
| 28_P_CW | *C. werkmanii* | 2 | C |
| 30_P_CF | *C. freundii* | 3 | A, B |
| 32_P_CF | *C. freundii* | 4 | A, B, D |
| 34_P_CF | *C. freundii* | 4 | A, F, B',G, |
| 9_E_CF | *C. freundii* | 3 | A, B |
| 38_P_CF | *C. freundii* | 5 | A, B'', G', H |
| 23_P_PA | *P. aeruginosa* | 3 | A, E |
| 27_P_CF | *C. freundii* | 4 | A, B, I |
| 29_P_CF | *C. freundii* | 4 | A, B, J |
| 13_E_CF | *C. freundii* | 5 | A, B'', G', H |

**B)**

| Plasmid | Length | GC % | Res. Genes | Closest Hit (NCBI) | Query cover | Identity % |
|---|---|---|---|---|---|---|
| A | 39882 | 58,92 | yes | LT837805.1 | 63% | 97 |
| A' | 39371 | 58,82 | yes | LT837805.2 | 64% | 97 |
| B | 88318 | 45,02 | no | CP033074.1 | 39% | 78 |
| B' | 86220 | 45,09 | no | CP033074.2 | 37% | 78 |
| B'' | 85137 | 45,18 | no | CP033074.2 | 38% | 78 |
| C | 163147 | 51,8 | yes | CP011977.1 | 18% | 99 |
| D | 89139 | 52,09 | no | CP024881.1 | 98% | 99 |
| E | 7173 | 56,6 | no | CP027172.1 | 94% | 99 |
| F | 2009 | 63,27 | no | LN853727.1 | 14% | 75 |
| G | 86638 | 49,21 | no | CP023978.0 | 5% | 98 |
| G' | 85406 | 49,24 | no | CP023978.1 | 4% | 98 |
| H | 6431 | 39,08 | no | AY842156.1 | 4% | 86 |
| I | 76389 | 51,84 | no | CP016762.1 | 79% | 99 |
| J | 162283 | 49,42 | no | CP024882.1 | 73% | 99 |

**Table S4.** Overview of all plasmids identified in the samples. **Table S4A** shows all samples and their respective plasmids, assembled using the hybrid Nanopore/Illumina assembly tool UniCycler. **Table S4B** shows detailed information about all identified plasmids. Plasmids A, B and C are involved in the resistance gene transmission and plasmid fusion and are discussed in detail in the main manuscript. Plasmids originating from the same ancestor are labelled with the same letter and apostrophes indicating versions with minor variations.

| Sample ID | Missing resistance genes |
|---|---|
| 27_P_C | qacH |
| 10_E_PA | aph(3')-XV |
| 10_P_PA | qacH |
| 11_E_PA | aph(3')-XV |
| 12_E_PA | aph(3')-XV, qacH |
| 14_E_PA | qacH |
| 19_P_PA | qacH |
| 22_P_PA | qacH |
| 24_P_PA | qacH, blaOXA-10 |
| 25_P_PA | qacH, blaOXA-10 |
| 31_P_PA | qacH (partial) |
| 7_E_PA | aac(6')-lb-cr, qacH, aac(6')-lb-cr, blaOXA-10 |
| 8_E_PA | qacH, blaOXA-10 |

**Table S5**. List of the samples with deletions affecting one ore more resistance genes.

### Supplementary References


1. Milne I, Stephen G, Bayer M, Cock PJ, Pritchard L, Cardle L, Shaw PD, Marshall D: **Using Tablet for visual exploration of second-generation sequencing data.** *Brief Bioinform* 2013, **14:**193-202.
2. Willmann M, Bezdan D, Zapata L, Susak H, Vogel W, Schroppel K, Liese J, Weidenmaier C, Autenrieth IB, Ossowski S, Peter S: **Analysis of a long-term outbreak of XDR Pseudomonas aeruginosa: a molecular epidemiological study.** *J Antimicrob Chemother* 2015, **70:**1322-1330.
3. Peter S, Bezdan D, Oberhettinger P, Vogel W, Dorfel D, Dick J, Marschal M, Liese J, Weidenmaier C, Autenrieth I, et al: **Whole-genome sequencing enabling the detection of a colistin-resistant hypermutating Citrobacter werkmanii strain harbouring a novel metallo-beta-lactamase VIM-48.** *Int J Antimicrob Agents* 2018, **51:**867-874.
4. Peter S, Oberhettinger P, Schuele L, Dinkelacker A, Vogel W, Dorfel D, Bezdan D, Ossowski S, Marschal M, Liese J, Willmann M: **Genomic characterisation of clinical and environmental Pseudomonas putida group strains and determination of their role in the transfer of antimicrobial resistance genes to Pseudomonas aeruginosa.** *BMC Genomics* 2017, **18:**859.
5. Gurevich A, Saveliev V, Vyahhi N, Tesler G: **QUAST: quality assessment tool for genome assemblies.** *Bioinformatics* 2013, **29:**1072-1075.
